# Risk factors and immunological characteristics of SARS-CoV-2 RNA re-positivity in convalescent COVID-19 Patients: A retrospective study

**DOI:** 10.64898/2026.09.11.750839

**Authors:** Li Li, Xin Zhang, Jingtong Fan, Jiaran Fan, Junxiao Du, Huimin Yan, Erhei Dai, Huixia Gao, Aidong Feng

**Affiliations:** Department of Emergency Medicine, The Fifth Hospital of Shijiazhuang, Shijiazhuang, China; Hebei Key Laboratory of Immune Mechanism of Major Infectious Diseases and New Technology of Diagnosis and Treatment, The Fifth Hospital of Shijiazhuang, Shijiazhuang, China; Department of Tuberculosis, The Fifth Hospital of Shijiazhuang, Shijiazhuang, China; Department of Medical Affairs, The Fifth Hospital of Shijiazhuang, Shijiazhuang, China

**Author notes:** Corresponding author: (AF). These authors contributed equally to this work (Li Li, Xin Zhang, Jingtong Fan, Jiaran Fan, Junxiao Du).

**Keywords:** SARS-CoV-2, COVID-19, Re-positivity, Risk factors, A/G ratio (albumin/globulin ratio), Immunological A (IgA)

## Abstract

**Purpose:** To investigate the incidence, clinical features, and risk factors for SARS-CoV-2 nucleic acid re-positivity in recovered COVID-19 patients.

**Methods:** We enrolled 441 convalescent COVID-19 patients from the rehabilitation ward of the Fifth Hospital of Shijiazhuang between January and April 2021. Based on nucleic acid test results, patients were categorized into repositive (n = 69) and non-repositive (n = 372) groups. Demographic, clinical, laboratory, and imaging data were compared between groups. Logistic regression was used to identify factors associated with re-positivity.

**Results:** The overall re-positivity rate was 15.65%. Compared with the non-repositive group, the repositive group presented with a significantly higher proportion of moderate disease severity (72.46% vs. 50.54%) and a lower proportion of patients under 14 years old (7.25% vs. 23.66%). Additionally, multi-lobar involvement (≥ 3 lobes) during the acute phase was more common in the repositive group (43.48% vs. 26.61%), with all differences being statistically significant (P < 0.001). Laboratory analysis revealed that repositive patients had significantly lower lymphocyte and eosinophil counts, as well as reduced albumin (ALB), aspartate aminotransferase/alanine aminotransferase (AST/ALT) ratio, and albumin/globulin (A/G) ratio. Furthermore, circulating CD3^+^, CD4^+^, and CD8^+^ T-cell counts were markedly decreased. In contrast, the neutrophil-to-lymphocyte ratio (NLR), immunoglobulin M (IgM), immunoglobulin A (IgA), and neutralizing antibody (NAb) levels were significantly elevated in re-positive patients (all P < 0.05). Multivariate logistic regression analysis identified the CD8^+^ as independent risk factors for re-positivity.

**Conclusion:** In convalescent patients, SARS-CoV-2 RNA re-positivity correlates with distinct clinical and immunological characteristics. The CD8^+^ T-cell level is an independent predictor, highlighting that dysregulated immune and inflammatory responses contribute to delayed viral clearance.

## Introduction

The global pandemic of COVID-19, caused by the highly transmissible RNA virus SARS-CoV-2, continues to present serious public health challenges. While the virus primarily targets the respiratory tract, it can also induce severe systemic complications, such as myocarditis and acute kidney injury, substantially increasing the overall disease burden [1]. In accordance with World Health Organization guidelines, patients are generally discharged following two consecutive negative reverse transcription-polymerase chain reaction (RT-PCR) tests [2]. However, during follow-up, some recovered individuals have tested positive again for viral RNA-a phenomenon commonly termed “re-positivity.” This phenomenon has raised significant public health concerns and highlights critical gaps in our understanding of viral persistence and clearance mechanisms [3].

Since the initial case series reported by Lan et al. in early 2020, repositive cases have been documented worldwide. Reported frequencies vary widely, ranging from 2.4% to 69.2%, with re-positivity observed from several days to months after initial recovery [4–6]. This substantial heterogeneity likely reflects differences in study populations, detection methodologies, viral variants, and follow-up protocols.

Available evidence suggests that repositive individuals generally exhibit lower viral loads and may have limited infectivity. However, their exact transmission risk and underlying immunological mechanisms remain poorly characterized [7]. Documented instances of intra-household transmission underscore the necessity for continued surveillance and preventive measures [8]. Thus, elucidating the clinical and immunological profile of re-positivity is essential to inform optimal patient management strategies.

Several mechanisms have been proposed to explain SARS-CoV-2 nucleic acid re-positivity, including persistent viral reservoirs, intermittent viral shedding, delayed immune clearance, and possible reinfection [9]. Among these, host immune competence, particularly the functional balance between humoral and cellular immunity, is widely recognized as a critical determinant of re-positivity risk [10]. To et al. documented a case of reinfection occurring 142 days after hospital discharge and suggested that waning humoral immunity against SARS-CoV-2 was a contributing factor to potential viral reactivation [11]. In a separate study, a 26-year-old patient with COVID-19 tested positive for SARS-CoV-2 again, with anti-SARS-CoV-2 antibodies detectable only after the onset of the second infection [12]. Although prior studies have provided valuable but fragmented insights, systematic assessments of comprehensive immunological markers and their associations with re-positivity in well-defined, homogeneous cohorts remain scarce.

To address this gap, we conducted a longitudinal analysis of 441 COVID-19 convalescents. We systematically compared clinical characteristics, laboratory parameters, and detailed immune profiles between repositive and non-repositive groups. Immune profiling included T-lymphocyte subsets quantified by flow cytometry, as well as SARS-CoV-2-specific immunoglobulin M (IgM), immunoglobulin G (IgG), immunoglobulin A (IgA), and neutralizing antibody (NAb) levels. This study aims to determine the incidence of re-positivity, identify independent clinical and immunological risk factors, and delineate associated immune signatures. Our findings are expected to provide a critical foundation for the early identification of high-risk individuals and for developing tailored post-recovery monitoring strategies.

## Methods

### Study design and participants

This study enrolled consecutive convalescent COVID-19 patients hospitalized in the Rehabilitation Ward of the Fifth Hospital of Shijiazhuang between January 2 and April 6, 2021. All participants were experiencing their initial infection with the SARS-CoV-2 and had not been vaccinated against COVID-19. Based on the presence or absence of a re-detectable positive SARS-CoV-2 RT-PCR result during the observation period, patients were categorized into two groups: a repositive group and a non-repositive group. The study protocol was approved by the Medical Ethics Committee of the Fifth Hospital of Shijiazhuang (Approval No. 2020008), and all participants provided written informed consent in accordance with the Declaration of Helsinki. The flow diagram of patient enrollment and grouping is shown in Fig 1.

**Fig 1.**
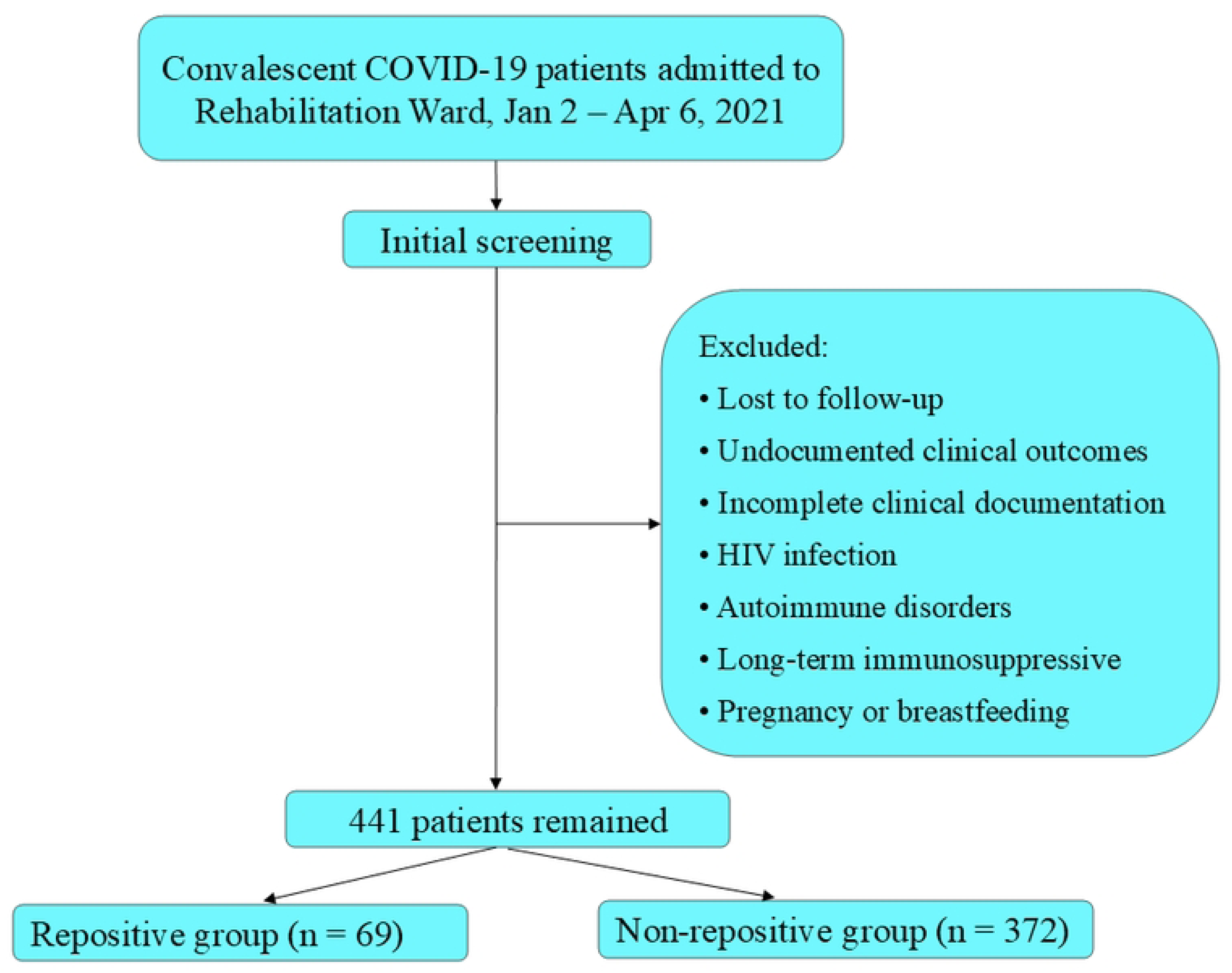
Flow diagram of patient enrollment, exclusion, and grouping in the retrospective cohort study of SARS-CoV-2 RNA re-positivity.

### Inclusion and exclusion criteria

Eligible participants were patients with confirmed COVID-19 who had been discharged as cured according to the Diagnosis and Treatment Protocol for Novel Coronavirus Pneumonia (Trial Version 8) [13]. The discharge criteria required: (i) normal body temperature for over three consecutive days; (ii) significant improvement in respiratory symptoms; (iii) chest imaging showing substantial resolution of acute exudative lesions; and (iv) two consecutive negative SARS-CoV-2 nucleic acid tests (sampled at least 24 hours apart). Additional requirements included provision of signed informed consent and availability of complete clinical records.

We excluded patients who were lost to follow-up, had undocumented clinical outcomes, or lacked complete clinical documentation. Further exclusions applied to individuals with HIV infection, autoimmune disorders, those receiving long-term immunosuppressive treatment (such as systemic corticosteroids), and pregnant or breastfeeding women. No missing data were observed for any primary or secondary outcome variables in the final study cohort.

### Data Collection and Clinical Sampling Protocol

Data were extracted from the electronic medical record system using a predefined case report form, which was completed by trained clinicians. The form captured demographic characteristics, underlying comorbidities (including cardiovascular diseases, diabetes, chronic respiratory diseases, and malignancies), clinical manifestations, treatment procedures, laboratory test results, SARS-CoV-2 antibody assays, and other relevant clinical information. To ensure accuracy, all entered data were verified independently by two data administrators.

Following the fulfillment of discharge criteria, patients were transferred to a rehabilitation ward for continued observation. Based on the follow-up protocol, an initial blood sample was drawn within 7 days, with subsequent samples collected at 7-day intervals. Nasopharyngeal swabs for nucleic acid testing were obtained at least twice per week. With respect to radiographic assessment, the initial chest X-ray or spiral computed tomography (CT) scan was acquired within 7 days of hospital admission. Follow-up imaging was then performed every 5 to 7 days based on clinical progression until discharge.

### Chemiluminescence immunoassay for SARS-CoV-2 antibodies

Anti-SARS-CoV-2 IgG, IgM, IgA, and NAb levels were quantitatively measured using a chemiluminescence immunoassay (iFlash 3000-C analyzer; YHLO Biotech Co., Ltd., Shenzhen, China). Serum samples were incubated with paramagnetic microparticles coated with SARS-CoV-2 antigens to facilitate antibody binding and immune complex formation. Following incubation, immune complexes were immobilized using a magnetic field, and nonspecific components were removed by washing. Chemiluminescence was subsequently initiated, and emitted light was quantified as relative light units (RLUs). All procedures were performed according to the manufacturer’s instructions. An antibody concentration ≥10 AU/mL was considered seropositive.

### Statistical analysis

All statistical analyses were performed using R software (version 4.1.3; R Foundation for Statistical Computing, Vienna, Austria). Continuous variables were assessed for normality using the Shapiro-Wilk test. Normally distributed data were expressed as mean ± standard deviation (SD) and compared between groups using independent samples t-tests or one-way analysis of variance (ANOVA). Non-normally distributed data were presented as median (interquartile range, IQR) and compared using the Wilcoxon rank-sum test. Categorical variables were summarized as frequencies and percentages, with group differences assessed using Pearson’s chi-square test or Fisher’s exact test, as appropriate.

Univariate logistic regression was performed to evaluate potential factors associated with nucleic acid re-positivity. Variables with *P* < 0.05 in univariate analysis were subsequently entered into a multivariate logistic regression model. Results are presented as odds ratios (ORs) with corresponding 95% confidence intervals (CIs) and *P* values. A two-sided P value < 0.05 was considered statistically significant for all analyses.

## Results

### Baseline characteristics and univariate comparisons

To identify factors associated with SARS-CoV-2 RNA re-positivity, we compared baseline demographic and clinical characteristics between repositive and non-repositive convalescent COVID-19 patients. Among 441 patients under medical observation, 69 (15.65%) experienced re-positivity. Age-stratified analysis revealed a significantly higher proportion of patients aged 14-59 years and a lower proportion of those aged <14 years in the repositive group (both P < 0.007). In terms of clinical severity, the majority of repositive cases were classified as moderate (72.46%), whereas asymptomatic and mild cases were less prevalent. In contrast, the non-repositive group had a higher proportion of asymptomatic infections (28.49%), with a statistically significant difference between the two groups (P = 0.001). No significant differences were observed in sex distribution or underlying comorbidities (P > 0.05) (Table 1).

**Table 1.** Demographic characteristics of convalescent COVID-19 patients.

| Characteristic | All patients<br>(n=441) | Repositive group<br>(n=69) | Non-repositive<br>group (n=372) | P value |
| --- | --- | --- | --- | --- |
| Sex |  |  |  | 0.080 |
| Male | 176 (39.90%) | 21 (30.43%) | 155 (41.67%) |  |
| Female | 265 (60.1%) | 48 (69.57%) | 217 (58.33%) |  |
| Age, years |  |  |  | 0.007 |
| <14 | 93 (21.09%) | 5 (7.25%) | 88 (23.66%) |  |
| 14-59 | 266(60.32%) | 47 (68.12%) | 219 (58.87%) |  |
| ≥60 | 82(18.59%) | 17 (24.63%) | 65(17.47%) |  |
| Clinical classification |  |  |  | 0.001 |
| Asymptomatic | 111 (25.17%) | 5 (7.25%) | 106 (28.49%) |  |
| Mild | 80 (18.14%) | 11 (15.94%) | 69 (18.55%) |  |
| Moderate | 238 (53.97%) | 50 (72.46%) | 188 (50.54%) |  |
| Severe | 12 (2.72%) | 3 (4.35%) | 9 (2.42%) |  |
| underlying disease | 120(27.21%) | 21(30.43%) | 99(26.61%) | 0.585 |
| Hypertension | 78(17.69%) | 13 (18.84%) | 65 (17.47%) | 0.689 |
| Chronic respiratory<br>disease | 14(3.17%) | 3 (4.34%) | 11 (2.96%) | 1.000 |
| Diabetes | 31(7.03%) | 7 (10.14%) | 24 (6.45%) | 0.353 |
| Cardiovascular disease | 27 (6.12%) | 6 (8.70%) | 21 (5.65%) | 0.439 |
| Cerebrovascular disease | 8 (1.81%) | 1 (1.45%) | 7 (1.88%) | 1.000 |
| Chronic liver disease | 11 (2.49%) | 2 (2.90%) | 9 (2.42%) | 1.000 |
| Malignant tumor | 4 (0.91%) | 1 (1.45%) | 3 (0.80%) | 0.484 |

### Clinical characteristics during the convalescent phase

To assess the association between clinical presentation and SARS-CoV-2 RNA re-positivity, we analyzed symptomatic and radiographic data from convalescent patients. Among the 441 patients included, most reported substantial symptomatic improvement during recovery, with only a minority experiencing residual symptoms.

The most common residual symptoms included cough and expectoration (10.88%), followed by headache or dizziness (2.72%), bloating (2.72%), chest tightness or dyspnea (2.49%), nausea or vomiting (1.59%), and fatigue (1.13%). However, none of these symptoms showed statistically significant differences between the repositive and non-repositive groups (all P > 0.05).

In contrast, radiographic findings revealed significant intergroup differences. Specifically, chest CT demonstrated that the extent of pulmonary lobe involvement was significantly greater in the repositive group than in the non-repositive group. Notably, this difference was observed both during the acute phase and at follow-up during recovery (both P < 0.001). With respect to treatment, the proportion of patients receiving antiviral therapy did not differ significantly between the two groups (26.09% vs. 23.39%, P > 0.05). Following medical observation, all patients recovered and were discharged without fatalities (Table 2).

**Table 2.** Clinical symptoms and imaging data of convalescent COVID-19 patients.

| Classification | All patients<br>(n=441) | Repositive group<br>(n=69) | Non-repositive group<br>(n=372) | *P* value |
| --- | --- | --- | --- | --- |
| Clinical symptoms during the recovery period |  |  |  |  |
| Fatigue | 5 (1.13%) | 1 (1.45%) | 4 (1.08%) | 0.563 |
| Cough, expectoration | 48(10.88%) | 8(11.59%) | 40(10.75%) | 0.763 |
| Chest tightness, Dyspnea | 11(2.49%) | 1(1.45%) | 10(2.69%) | 0.884 |
| Headache, dizziness | 12(2.72%) | 3(4.35%) | 9(2.42%) | 1.000 |
| Nausea, vomiting | 7(1.59%) | 3(4.35%) | 4(1.08%) | 0.127 |
| Bloating | 12(2.72%) | 1(1.45%) | 11(2.96%) | 0.792 |
| Acute-phase chest CT |  |  |  | <0.001 |
| No abnormalities | 159(36.05%) | 9(13.04%) | 150(40.32%) |  |
| Involvement of a single lung lobe | 91(20.63%) | 12(17.39%) | 79(21.24%) |  |
| Involvement of two lung lobes | 62(14.06%) | 18(26.09%) | 44(11.83%) |  |
| Involvement of three lung lobes | 129(29.25%) | 30(43.48%) | 99(26.61%) |  |
| Convalescent-phase chest CT |  |  |  | <0.001 |
| No abnormalities | 248(56.24%) | 22(31.88%) | 226(60.75%) |  |
| Involvement of a single lung lobe | 74(16.78%) | 20(28.99%) | 54(14.52%) |  |
| Involvement of two lung lobes | 57(12.93%) | 11(15.94%) | 46(12.37%) |  |
| Involvement of three lung lobes | 62(14.06%) | 16(23.19%) | 46(12.37%) |  |
| Antiviral therapy | 105(23.81%) | 18(26.09%) | 87(23.39%) | 0.685 |
| Outcome |  |  |  |  |
| Recovered and discharged | 441(100%) | 69(100%) | 372(100%) |  |
| Death | 0 | 0 | 0 |  |

### Laboratory profiles and immune responses

#### Hematological and biochemical parameters

Although COVID-19 predominantly presents with respiratory symptoms, growing evidence highlights its multisystemic involvement [14,15]. To evaluate whether SARS-CoV-2 RNA re-positivity during convalescence is associated with systemic physiological disturbances, we compared hematological and biochemical profiles between repositive and non-repositive patients.

As shown in Table 3, the repositive group exhibited significantly lower lower lymphocyte counts (P = 0.014), eosinophil counts (P = 0.004), ALB levels (P = 0.009), AST/ALT ratios (P = 0.010), and A/G ratios (P = 0.034) compared with the non-repositive group. Conversely, the neutrophil-to-lymphocyte ratio (NLR) (P = 0.003) levels were significantly elevated in repositive patients. No significant differences were observed in other laboratory parameters between the two groups.

**Table 3.** Comparison of Hematological Parameters in COVID-19 Patients.

| Classification | All patients(n = 441) | Repositive group<br>(n = 69) | Non-repositive group<br>(n = 372) | P value |
| --- | --- | --- | --- | --- |
| WBC (*10<sup>9</sup>/L) | 6.38 (5.33, 7.44) | 6.43 (5.33, 7.73) | 6.36 (5.33, 7.42) | 0.692 |
| NEUT (*10<sup>9</sup>/L) | 3.61 (2.88, 4.46) | 3.84 (3.05, 4.88) | 3.58 (2.88, 4.31) | 0.052 |
| LYMPH<br>(*10<sup>9</sup>/L) | 2.06 (1.71, 2.54) | 1.99 (1.54, 2.34) | 2.06 (1.74, 2.61) | 0.014 |
| MONO (*10<sup>9</sup>/L) | 0.35 (0.29, 0.43) | 0.36 (0.29, 0.42) | 0.35 (0.29, 0.43) | 0.762 |
| EOS (*10<sup>9</sup>/L) | 0.13 (0.08, 0.20) | 0.09 (0.06, 0.16) | 0.13 (0.09, 0.21) | 0.004 |
| RBC (*10<sup>12</sup>/L) | 4.45 (4.16, 4.84) | 4.39 (4.08, 4.72) | 4.47 (4.17, 4.86) | 0.182 |
| PLT(*10<sup>9</sup>/L) | 254.00<br>(209.00, 290.00) | 250.50<br>(206.25, 285.75) | 256.50<br>(209.75, 295.25) | 0.372 |
| NLR | 1.72 (1.36, 2.34) | 2.07 (1.46, 2.72) | 1.67 (1.34, 2.25) | 0.003 |
| PLR | 117.99<br>(96.11, 150.31) | 121.97<br>(99.30, 159.66) | 116.47<br>(96.01, 148.96) | 0.203 |
| ALT (U/L) | 20.60 (15.08, 31.73) | 24.85 (16.90, 38.28) | 20.10 (14.93, 30.20) | 0.049 |
| AST (U/L) | 19.20 (16.58, 25.48) | 19.50 (16.10, 24.70) | 19.20 (16.60, 25.70) | 0.802 |
| ALB (g/L) | 44.80 (42.50, 47.10) | 43.80 (41.28, 46.30) | 44.90 (42.70, 47.20) | 0.009 |
| GLB (g/L) | 24.00 (21.80, 27.00) | 24.30 (22.25, 27.35) | 23.80 (21.75, 26.95) | 0.165 |
| LDH (U/L) | 166.00<br>(148.00, 193.00) | 166.00<br>(147.50, 187.25) | 166.00<br>(148.00, 193.00) | 0.704 |
| ALP (U/L) | 71.80 (57.08, 95.63) | 71.10 (57.35, 93.10) | 72.00 (56.85, 97.85) | 0.915 |
| GGT (U/L) | 17.30 (11.90, 28.15) | 20.6 (13.25, 27.55) | 16.70 (11.50, 28.55) | 0.087 |
| GLU (mmol/L) | 5.31 (4.90, 5.73) | 5.27 (4.84, 5.68) | 5.32 (4.93, 5.74) | 0.429 |
| BUN (mmol/L) | 5.04 (4.21, 5.96) | 5.14 (4.33, 6.16) | 5.03 (4.16, 5.89) | 0.474 |
| Cr (umol/L) | 64.90 (55.30, 75.83) | 66.10 (57.45, 75.58) | 64.70 (54.55, 75.98) | 0.181 |
| AST/ALT | 0.88 (0.69, 1.21) | 0.76 (0.62, 1.00) | 0.93 (0.70, 1.25) | 0.010 |
| A/G | 1.84 (1.62, 2.09) | 1.77 (1.58, 1.99) | 1.86 (1.63, 2.09) | 0.034 |

#### Analysis of T-cell subsets in convalescent patients

To assess immune status following SARS-CoV-2 infection, we analyzed T-lymphocyte subsets in 235 convalescent patients, including 34 repositive and 201 non-repositive cases. Compared with the non-repositive group, the repositive group showed significantly lower absolute counts of CD3^+^, CD4^+^, and CD8^+^ T cells (all P < 0.05). In contrast, the CD4^+^/CD8^+^ ratio did not differ significantly between the two groups (P > 0.05) (Fig 2).

**Fig 2.**
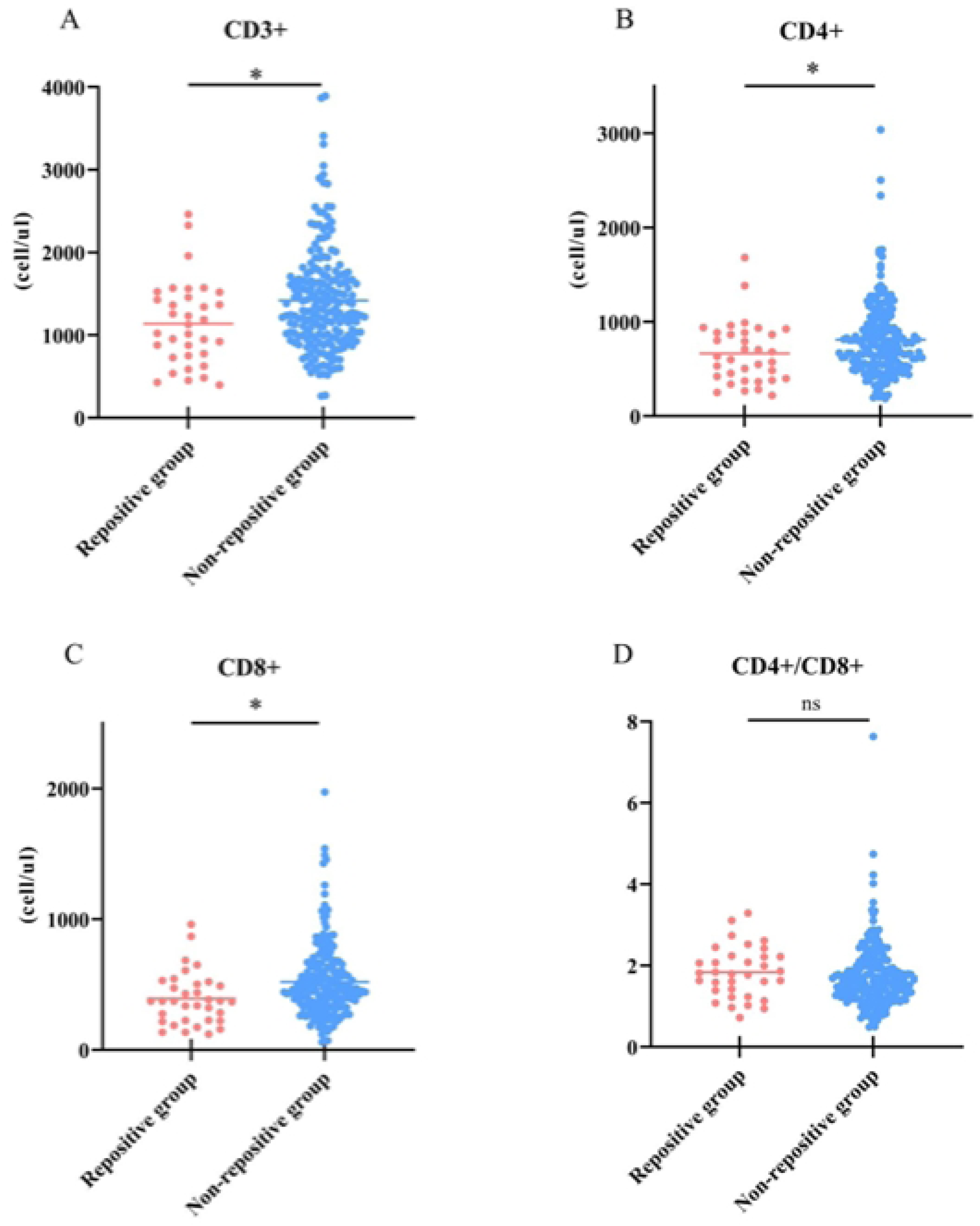
Comparison of T lymphocyte counts in repositive group and non-repositive group. Signiffcance levels were denoted as *P < 0.05, **P < 0.01, ***P < 0.001, and ****P < 0.0001.

#### Humoral immune responses in convalescent patients

Given the critical role of adaptive immunity in viral clearance and recovery, we compared humoral immune responses between repositive and non-repositive convalescent patients by measuring SARS-CoV-2-specific immunoglobulins and neutralizing antibodies. As summarized in Fig 3, levels of virus-specific IgM, IgA, and NAb were all significantly higher in the repositive group than in the non-repositive group (all P < 0.05). In contrast, IgG levels did not differ significantly between the two groups (P > 0.05).

**Fig 3.**
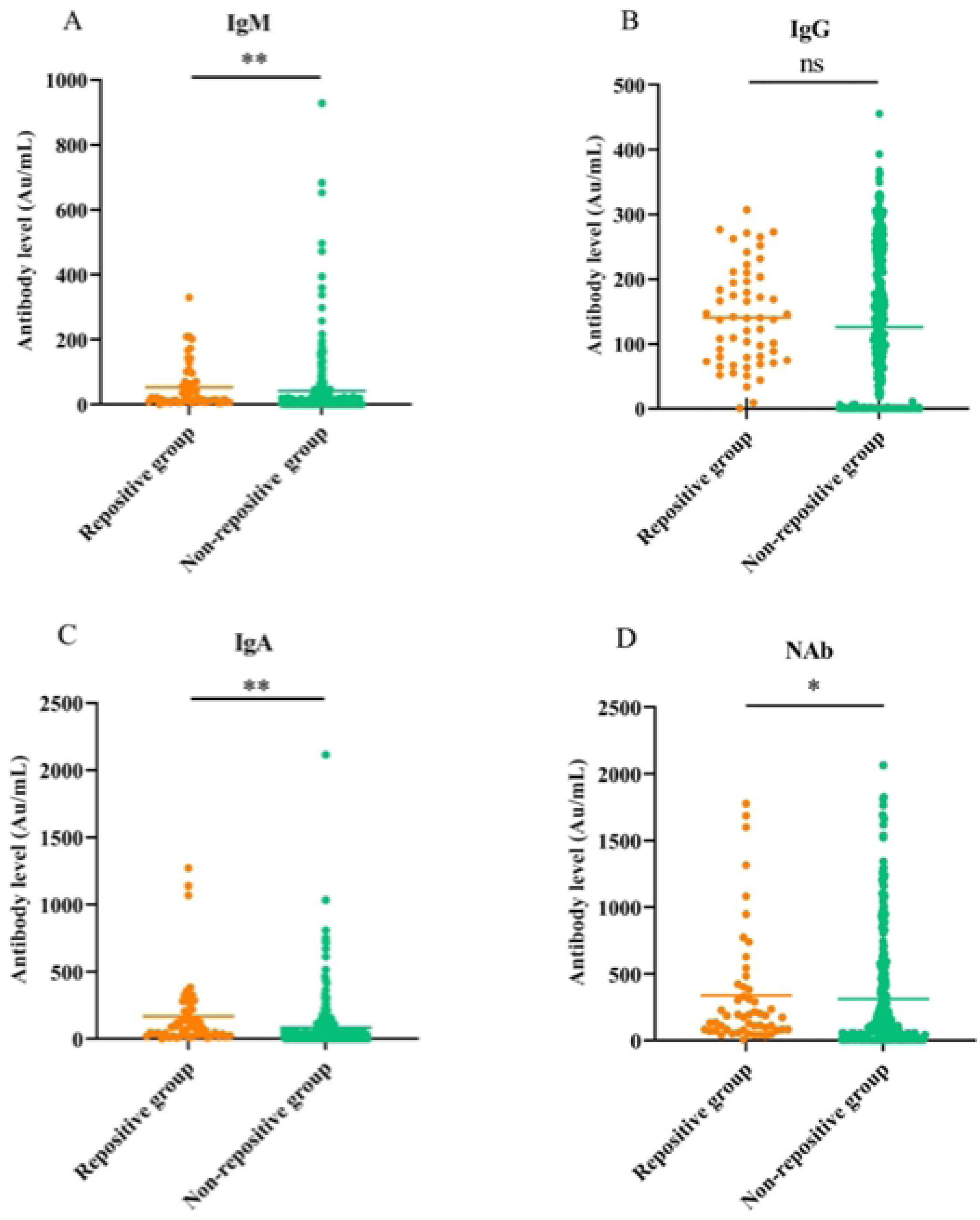
Comparison of SARS-CoV-2-specific antibodies in repositive group and non-repositive group. Signiffcance levels were denoted as *P < 0.05, **P < 0.01, ***P < 0.001, and ****P < 0.0001.

#### Independent predictors of re-positivity

Based on the comparative analyses described above, significant intergroup differences were identified between repositive and non-repositive patients. These differences spanned multiple domains, including demographic characteristics (age, clinical classification), radiographic findings (chest CT manifestations), hematological parameters (LYMPH, NLR), biochemical markers (AST/ALT ratio, ALB, GLB, A/G ratio), immunological indices (CD8^+^ T-cell counts), and serological markers (IgA). To identify independent factors associated with re-positivity, a multivariate logistic regression analysis was subsequently performed. The results demonstrated that the CD8^+^ T-cell level was an independent predictor of re-positivity (P < 0.05; Tables 4 and 5).

**Table 4.** Univariate logistic regression analysis of recurrent PCR positivity in convalescent COVID-19 patient.

| Variable | Estimate | Std.Error | OR( 95%CI) | *P* value |
| --- | --- | --- | --- | --- |
| Age | 0.019 | 0.006 | 1.019(1.006-1.032 ) | 0.004 |
| Disease classification | 0.072 | 0.183 | 2.057(1.436-2.946) | <0.001 |
| LYMPH | -0.499 | 0.212 | 0.607(0.400-0.920) | 0.019 |
| EO | -2.618 | 1.421 | 0.073(0.004-1.183) | 0.065 |
| NLR | 0.512 | 0.152 | 1.699(1.239-2.248) | <0.001 |
| AST/ALT | -0.901 | 0.388 | 0.406(0.190-0.869) | 0.020 |
| ALB | -0.079 | 0.035 | 0.924(0.862-0.990) | 0.025 |
| A/G | -0.843 | 0.035 | 0.431(0.197-0.942) | 0.035 |
| CD3<sup>+</sup> T | -0.001 | 0.000 | 0.999(0.998-1.000) | 0.15 |
| CD4<sup>+</sup> T | -0.001 | 0.001 | 0.999(0.997-1.000) | 0.41 |
| CD8<sup>+</sup> T | -0.003 | 0.001 | 0.997(0.995-0.999) | 0.010 |
| IgA | 0.002 | 0.001 | 1.002(1.000-1.003) | 0.010 |

**Table 5.** Multivariate Logistic Regression Analysis of Risk Factors for Recurrent SARS-CoV-2 RNA Positivity in Convalescent COVID-19 Patients.

| Variable | Estimate | Std.Error | OR(95%CI) | P value |
| --- | --- | --- | --- | --- |
| IgA | 0.001 | 0.001 | 1.001(1.000-1.003) | 0.062 |
| CD8<sup>+</sup> | -0.493 | 0.150 | 0.611(0.455-0.810) | 0.001 |

Lower CD8^+^ T-cell count was identified as an independent risk factor for SARS-CoV-2 re-positivity by multivariate logistic regression (OR = 0.611 per 100 cells/μL increase, 95% CI: 0.455–0.810, P = 0.001). Stratified analysis using the median cutoff of 480 cells/μL showed that patients with CD8^+^ T-cell counts below this threshold had an 18.1% absolute risk of re-positivity during the 14-day follow-up period, nearly double the 9.8% risk observed in patients with higher counts (RR = 1.84, 95% CI: 0.92–3.68). The absolute risk difference was 8.3%, indicating that approximately 12 patients with low CD8^+^ T-cell counts would need to be closely monitored to detect one additional case of re-positivity.

## Discussion

COVID-19, representing the third coronavirus pandemic of the 21st century following SARS and MERS, remains a critical and ongoing global public health threat. A notable proportion of convalescent patients experience subsequent recurrent positivity for SARS-CoV-2 RNA upon retesting. In this context, the present study was conducted to provide a comprehensive characterization of this re-positivity phenomenon among recovered COVID-19 patients.

In our cohort, the re-positivity rate for SARS-CoV-2 nucleic acid was 15.65%, which falls within the range reported in earlier studies [16,17]. Advanced age was significantly associated with an increased likelihood of re-positivity. Clinically, repositive cases were predominantly of moderate severity and exhibited more extensive pulmonary involvement on imaging. Laboratory analysis further revealed a distinct profile in these patients, marked by lower levels of nutritional and immune competence markers, such as ALB, together with elevated indicators of systemic inflammation and humoral immune response.

Regarding high-risk populations, our data suggest that age is an influential factor, with children (<14 years) showing a relatively lower risk. This observation contrasts with reports highlighting a higher risk among adolescents or initially mild-symptomatic individuals [18]. The proportion of patients aged ≥60 years was significantly higher in the repositive group, which may be attributed to the diminished humoral immune response associated with age-related immune cell senescence and the expansion of a pro-inflammatory B-cell subset in the elderly [19].

Mechanistically, an elevated NLR is an established marker of adverse outcomes in acute COVID-19 [20]. While some studies associate higher pre-discharge lymphocyte counts with accelerated viral clearance [21]. others identify lymphocytosis as a potential risk factor for disease recurrence [22]. In contrast, our data demonstrate significantly lower lymphocyte counts in the re-positivity group, suggesting distinct patterns of immune reconstitution following infection.

Analysis of T-cell subsets showed significantly lower levels of CD3^+^, CD4^+^, and CD8^+^ T cells in repositive patients compared to controls. Univariate analysis indicated that reduced T-cell counts may impede effective viral clearance, thereby increasing re-positivity risk. This observation aligns with findings reported by Yao et al., who also described an association between low CD4^+^ T-cell levels and repositive outcomes [16]. Accumulating evidence suggests that T-cell exhaustion and functional impairment likely contribute to inadequate immune responses, promoting viral re-positivity [23].

Notably, this study identified a concomitant presentation of a reduced A/G ratio and elevated serum IgA levels in the context of chronic inflammation. The diminished A/G ratio, principally attributable to hypoalbuminemia, is a recognized indicator of a persistent hypercatabolic state and systemic nutritional depletion. This metabolic alteration signifies the mobilization of endogenous protein reserves to sustain chronic inflammatory processes, thereby undermining the substrate availability crucial for competent immune cell function. Conversely, the elevated IgA level implies a distinct trajectory of immune dysregulation. This finding likely denotes aberrant B-cell activation rather than a protective mucosal immune response. Such activation may be precipitated by continuous antigenic stimulation, possibly arising from inflammation-induced barrier compromise, culminating in a non-resolving immune reaction with the potential to inflict tissue damage [24–25].

Based on the observational and immunological data from this study, we propose a clinically anchored immunometabolic axis imbalance model to elucidate the mechanism underlying SARS-CoV-2 RNA re-positivity in convalescent COVID-19 patients (Fig 4). Patients who developed subsequent re-positivity exhibited baseline high-risk features: older age, moderate disease severity, and multi-lobar (≥3 lobes) pulmonary involvement on chest CT. These features correlated with persistent, synergistic dysregulation across three interconnected domains, nutritional-metabolic dysfunction, impaired cellular immunity, and aberrant humoral activation, which collectively form a self-reinforcing pathological axis that impedes residual viral clearance [26].

**Fig 4.**
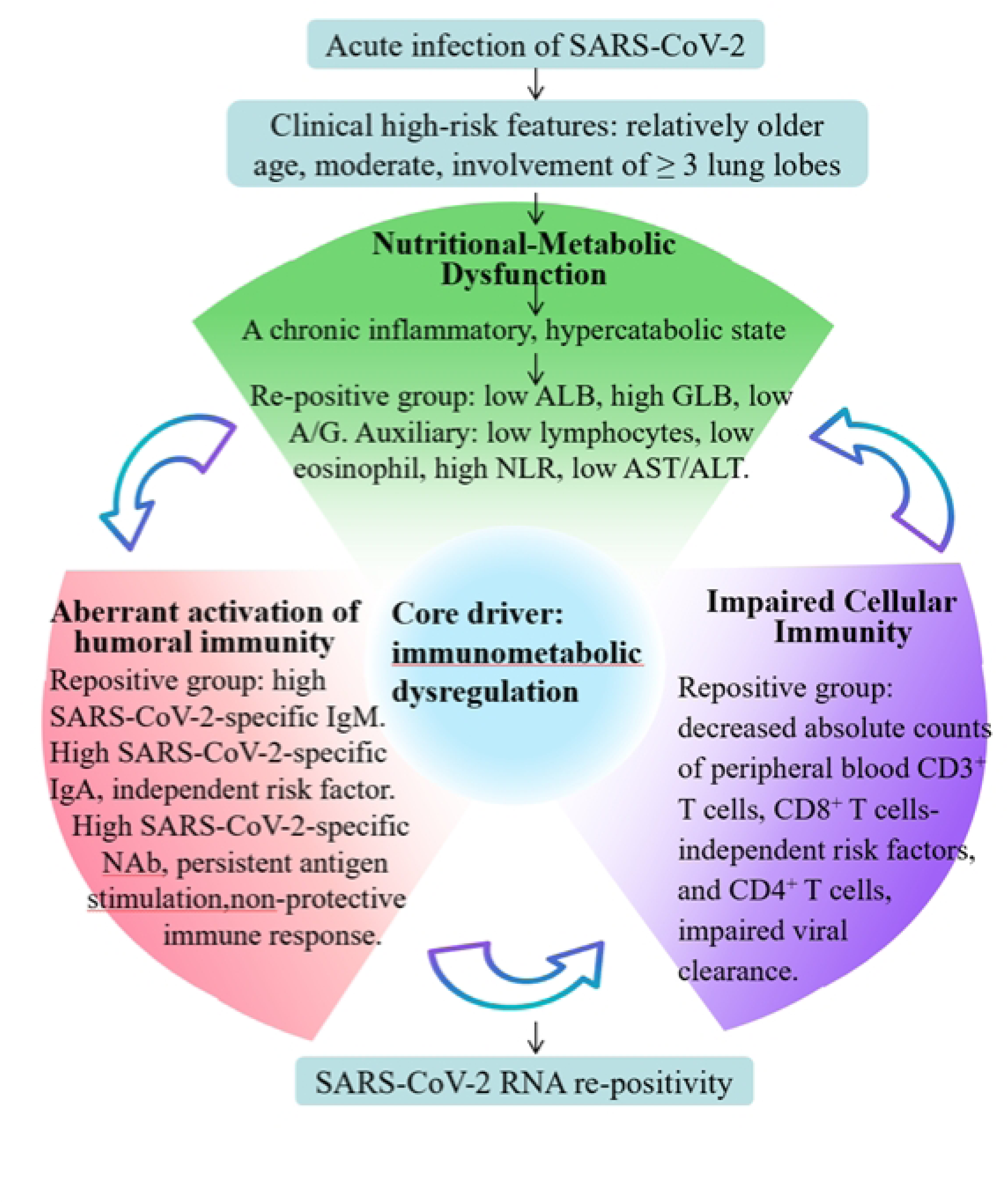
Imbalance in the immune-metabolic axis explains SARS-CoV-2 re-positivity.

(1) Nutritional-Metabolic Dysfunction

Manifested as persistent hypercatabolic inflammation: markedly reduced ALB, elevated GLB, and consequently a decreased A/G ratio. Concomitant abnormalities included lymphopenia, eosinopenia, elevated NLR, and reduced AST/ALT ratio. These findings indicate systemic nutritional depletion and unresolved low-grade inflammation, which compromise the metabolic substrate supply critical for effective antiviral immunity [27].

(2) Impaired Cellular Immunity

Repositive patients had significantly reduced absolute counts of CD3⁺, CD4⁺, and CD8⁺ T lymphocytes (an independent predictor of re-positivity), indicating attenuated T-cell-mediated immune surveillance. This directly impairs the capacity to eliminate residual viral RNA, thereby increasing susceptibility to re-positivity during convalescence. Studies have shown reduced T-cell responses in COVID-19 survivors even months after infection, raising concerns about diminishing immunity and potential reinfection [28].

(3) Aberrant activation of humoral immunity

The convalescent cohort exhibited markedly elevated levels of SARS-CoV-2-specific IgM, IgA, and neutralizing antibodies (NAbs), whereas no statistically significant difference in IgG levels was observed between groups. Neutralizing antibody levels measured in the early disease phase have also been identified as a correlate of risk for SARS-CoV-2 RNA re-positivity [29]. A disproportionate elevation of IgM and IgA relative to protective IgG responses points to chronic immune activation sustained by viral persistence, rather than robust, productive viral clearance[30]. These findings align with the concept that long COVID involves immune dysregulation, wherein the immune system remains hypervigilant, accompanied by persistent inflammation [31].

Dynamic crosstalk between these three components establishes a self-reinforcing vicious cycle: nutritional-metabolic dysfunction impairs T-cell function → impaired cellular immunity abrogates B-cell regulation, driving dysregulated humoral activation → persistent inflammation further exacerbates metabolic depletion. Ultimately, this immunometabolic axis imbalance prevents complete clearance of residual SARS-CoV-2 RNA, manifesting clinically as recurrent re-positivity following recovery.

Several limitations must be acknowledged. First, as a single-center retrospective study, the generalizability of our findings is limited. Second, systematic follow-up data on long-term outcomes such as long COVID in patients with recurrent PCR positivity are lacking. Finally, viral culture was not performed to confirm the infectivity of repositive samples. Despite these limitations, the study has notable strengths, particularly the high homogeneity of the cohort all participants were infected with SARS-CoV-2, were unvaccinated, and were experiencing their first infection which helped minimize confounding.

## Conclusion

In summary, this study systematically delineates the clinical features and underlying immunometabolic disturbances in a highly homogeneous cohort of convalescent COVID-19 patients with recurrent SARS-CoV-2 nucleic acid positivity. We identify significant associations between disease severity and the recurrence of positivity, thereby providing potential biomarker signatures for identifying high-risk individuals during follow-up. The proposed “immune–metabolic axis” imbalance model advances the mechanistic understanding of post-infection viral persistence and offers a novel theoretical framework for investigating the long-term evolution of immune responses following viral clearance.

## Acknowledgements

We wish to thank all the participants for taking part in the study. We also thank the Fifth Hospital of Shijiazhuang for its assistance in the implementation of the project. The authors declare no conflicts of interest.

## Author Contributions

**Conceptualization:** Li Li, Xin Zhang,

**Data curation:** Li Li, Xin Zhang, Meng Meng, Jingtong Fan, Junxiao Du.

**Formal analysis:** Li Li, Xin Zhang, Qiao Liu, Jiaran Fan, Aidong Feng, Huixia Gao.

**Investigation:** Li Li, Xin Zhang, Huimin Yan, Aidong Feng.

**Methodology:** Li Li, Xin Zhang, Erhei Dai, Aidong Feng, Huixia Gao.

**Project administration:** Li Li, Xin Zhang,

**Software:** Li Li, Xin Zhang,

**Supervision:** Li Li, Xin Zhang, Aidong Feng.

**Validation:** Li Li, Xin Zhang, Aidong Feng.

**Visualization:** Li Li, Xin Zhang, Junxiao Du.

**Writing– original draft:** Li Li, Xin Zhang, Jingtong Fan, Aidong Feng.

**Writing– review & editing:** Li Li, Xin Zhang, Jiaran Fan, Erhei Dai, Aidong Feng.

